# The molecular taxonomy of primate amygdala via single-nucleus RNA-sequencing analysis

**DOI:** 10.1101/2020.07.29.226829

**Authors:** Lei Zhang, Yanyong Cheng, Shihao Wu, Yufeng Lu, Zhenyu Xue, Dai Chen, Bo Zhang, Zilong Qiu, Hong Jiang

## Abstract

Amygdala is the central brain region governing emotional responses of mammals. However, the detailed molecular taxonomy of amygdala in higher mammals such as primates is still absent. Here we present the molecular taxonomy of amygdala in rhesus monkeys by single-cell RNA sequencing analysis. We found that there are five major cell types in primate amygdala, glutamatergic neurons, GABAergic neurons, astroglia, oligodendrocytes and oligodendrocyte progenitor cells (OPCs). Glutamatergic neurons in the primate amygdala exhibits a great diversity comparing to them in the rodent amygdala. In particular, GABAergic neurons in primate amygdala appeared to be quite unique and contain different cellular composition as them in rodent amygdala. The astroglia in primate amygdala contains two more subtypes comparing to astroglia in rodent amygdala. Taken together, although the evolutionary conservation, the molecular taxonomy study of primate amygdala provides critical insights for understanding the cellular architecture of the primate brain.

## INTRODUCTION

Amygdala is an evolutionally conserved brain structure across vertebrate animals. The physiological functions of amygdala were initially found to be related to emotionally responses to negative stimuli, such as fear and aversion^[1]^. However, accumulated evidences over the last a few decades strongly suggested that amygdala is involved in a wide range of behavioral responses including social, aggressive and parenting behaviors^[2, 3]^. Recent neural circuitry studies have revealed that amygdala is intensively connected with numerous brain regions, thus suggesting that amygdala plays a pivotal role in regulating animal adaptive behaviors. Specifically, dysfunctions of amygdala in human were associated with psychiatric and emotional disorders^[4, 5]^.

Although intensive works were carried out to analyze the role of amygdala in rodents, the anatomical structure of amygdala has changed enormously from rodents to primates after tens of million years of evolution^[2]^. Structurally, amygdala is composed of distinct nuclear components, including central amygdala (CeA), medial amygdala (MeA), as well as basal and lateral amygdala (BLA)^[6]^. Unlike rodents, the BLA nucleus possesses the majority region in primate brain, comparing to CeA and MeA^[2, 7]^. Since the cellular compositions varies significantly, across different amygdala nucleus, it is reasonable to deduce that the function of amygdala may vary in rodents and primates. Although the developmental and regional gene profiling in rhesus monkeys were performed previously, the gene expression information in single cell resolution is still not available^[8]^

Therefore, obtaining the comprehensive molecular taxonomy of amygdala would be a critical milestone, in order to fully understand the role of amygdala in the brain. Till now, the molecular taxonomy analysis of amygdala mainly focused on the MeA region of mouse amygdala using single-cell sequencing technology^[3, 9]^. The comprehensive molecular taxonomy for the whole amygdala in rodents or primates is still not available. In this work, we analyzed 26,400 cells collected from amygdala of two male rhesus monkeys (Macaca mulatta) using single-nucleus RNA-sequencing and acquired the largest taxonomy dataset for primate amygdala. This work would provide a valuable dataset for deepening our understanding cellular composition of primate brain.

## MATERIAL AND METHODS

### Collection of amygdala from rhesus macaque

The animal studies were performed according to the guidelines and regulations of the Institute of Laboratory Animal Science, Peking Union Medical College and Chinese Academy of Medical Science (Beijing, China). The use of rhesus macaque in research at the Institute of Laboratory Animal Science was approved by the Institutional Animal Care and Use Committee (Protocol number #XC17001). Efforts were made to minimize the number of animals in the studies. The rhesus macaques were purchased from the Institute of Laboratory Animal Science, Peking Union Medical College and Chinese Academy of Medical Science (Beijing, China). Two male rhesus macaques were used in the studies. The rhesus macaques received 2.5%–3% sevoflurane and 100% oxygen for the harvest of the amygdalae on P35.

### Nucleus Isolation

The tissues samples of amygdala were surgically removed and the frozen tissue was used to intact nucleus isolation. The nucleus was isolated and purified as previously described with some modifications^[10]^. Briefly, the frozen tissue was homogenized in NLB buffer which contain 250 mM Sucrose, 10 mM Tris-HCl, 3 mM MgAc2, 0.1% Triton X-100 (Sigma-Aldrich, USA), 0.1 mM EDTA, 0.2U/μL RNase Inhibitor (Takara, Japan). Various concentration of sucrose was used to purify the nucleus. The concentration of nucleus was adjusted to about 1000 nuclei / μL for snRNA-Seq.

### Single-cell RNA-Seq Experiment

The scRNA-Seq libraries were generated using the 10X Genomics Chromium Controller Instrument and Chromium Single Cell 3’ V2 Reagent Kits (10X Genomics, Pleasanton, USA). Briefly, cells nuclei were concentrated to 1000 nuclei/μL and approximately 15000 nuclei were loaded into each channel to generate single-cell Gel Bead-In-Emulsions (GEMs), which results into expected mRNA barcoding of 9000 single nucleus for each sample. After the RT step, GEMs were broken and barcoded-cDNA was purified and amplified. The amplified barcoded cDNA was fragmented, A-tailed, ligated with adaptors and index PCR amplified. The final libraries were quantified using the Qubit High Sensitivity DNA assay (Thermo Fisher Scientific, USA) and the size distribution of the libraries were determined using a High Sensitivity DNA chip on a Bioanalyzer 2200 (Agilent, USA). All libraries were sequenced by HiSeq Xten (Illumina, USA) on a 150 bp paired-end run.

### Single-nucleus RNA Sequencing Statistical Analysis

We applied fastp with default parameter filtering the adaptor sequence and removed the low-quality reads to achieve clean data. Feature-barcode matrices were obtained by aligning reads to the rhesus monkey genome (Version: Mmul 10, Ensembl) using CellRanger v3.0.0. In order to minimize the sample batch, we applied the down sample analysis among samples sequenced according to the mapped barcoded reads per cell of each sample and achieved the aggregated matrix eventually. Nucleus contained over 200 expressed genes and mitochondria UMI rate below 20% passed the quality filtering and mitochondria genes were removed in the expression table but used for nucleus expression regression to avoid the effect of the nucleus status for clustering analysis and marker analysis of each cluster.

Seurat package (version: 2.3.4, https://satijalab.org/seurat/) was used for nucleus normalization and regression based on the expression table according to the UMI counts of each sample and percent of mitochondria rate to obtain the scaled data. PCA was constructed based on the scaled data with all high variable genes and top 8 dims of 20 PCA dims were used for tSNE construction. Utilizing graph-based cluster method (resolution = 0.8), we acquired the unsupervised the nucleus cluster result and we calculated the marker genes by FindAllMarkers function with wilcox rank sum test algorithm under following criteria:1. logFC > 0.25; 2. P-value < 0.05; 3. min.pct > 0.1. Cell Type identification was applied based on the gene sets from the two brain scRNA-Seq articles^[11, 12]^. In order to identify the cell type detailed, several clusters was selected for re-tSNE analysis, graph-based clustering (20 dimension PCA with 8 dims used, resolution = 0.8) and marker analysis (wilcox rank sum test algorithm, 1. logFC > 0.25; 2. P-value < 0.05; 3. min.pct > 0.1) was applied based on the normalized data.

### Pseudo-Time Analysis

We applied the Single-Cell Trajectories analysis utilizing Monocle2 (http://cole-trapnell-lab.github.io/monocle-release) using DDR-Tree and default parameter. Before Monocle analysis, we select marker genes of the Seurat clustering result and raw expression counts of the cell passed filtering. Based on the pseudo-time analysis, branch expression analysis modelling (BEAM Analysis) was applied for branch fate determined gene analysis. We advise the start point in biological sense depended on known function marker genes.

### Pathway Analysis

Pathway analysis was used to find out the significant pathway of the differential genes according to KEGG database. The Fisher’s exact test was turned to select the significant pathway, and the threshold of significance was defined by P-value and FDR. The cases were selected when the calculated P□< □0.05^[13]^.

### LIGER analysis for macaque and mouse amygdala datasets

We used package LIGER1 (https://macoskolab.github.io/liger/) to integrate the two datasets by the function create Liger to evaluate the conversation and variation across species. We use the function toupper to convert all mouse gene names to uppercase. The function selects Genes (var. thresh = 0.1) was used to perform variable gene selection on macaque and mouse datasets separately and then takes the union. Next, we identify cells that load on corresponding cell factors and quantile normalize their factor loading across datasets. Cell dimensionality reduction was performed with function run tSNE. Function plot Gene Loadings was used to visualize the most highly loading genes (both shared and dataset specific) for each factor. To compare different cluster assignments, we employed the function makeriverplot to visualize the previously cell types assignments of macaque and mouse with liger joint clusters.

### Statistical analysis

Comparisons between two groups were made using *t*-tests. The quantification graphs were analysed by using GraphPad Prism (GraphPad Software).

## AVAILABILITY

CellRanger v3.0.0; Seurat package (version: 2.3.4, https://satijalab.org/seurat/); Monocle2 (http://cole-trapnell-lab.github.io/monocle-release); ggplot (http://had.co.nz/ggplot2/). R version 3.4.0.

## DATA AVAILABILITY

The sequencing data of single-cell sequencing has been uploaded to a publicly available database. [The raw data was uploaded to Gene Expression Omnibus (GEO) DataSets:145765]. (Web link: https://www.ncbi.nlm.nih.gov/geo/query/acc.cgi?acc=GSE145765).

## RESULTS

### Unbiased single-nuclear RNA sequencing reveals various cell types in primate amygdala

The comprehensive molecular and cellular taxonomy would rely on multiple information including morphology, electrophysiological property, and gene expression profiles^[14]^. Due to limited information in primate brains, in this work we primarily used profiling of gene expression as the major way to acquire the molecular taxonomy of amygdala.

Since primate brain tissues are normally heavily myelinated and intact cells are difficult to obtain during the dissociation process, we collected nucleus from amygdala tissues of two rhesus monkeys of postnatal 35 days old and performed 10XGenomics-based single-nuclear RNA sequencing (snRNA-seq) repetitively for three times with combined samples from two monkeys (Fig. 1A, Table S1).

**Figure 1.**
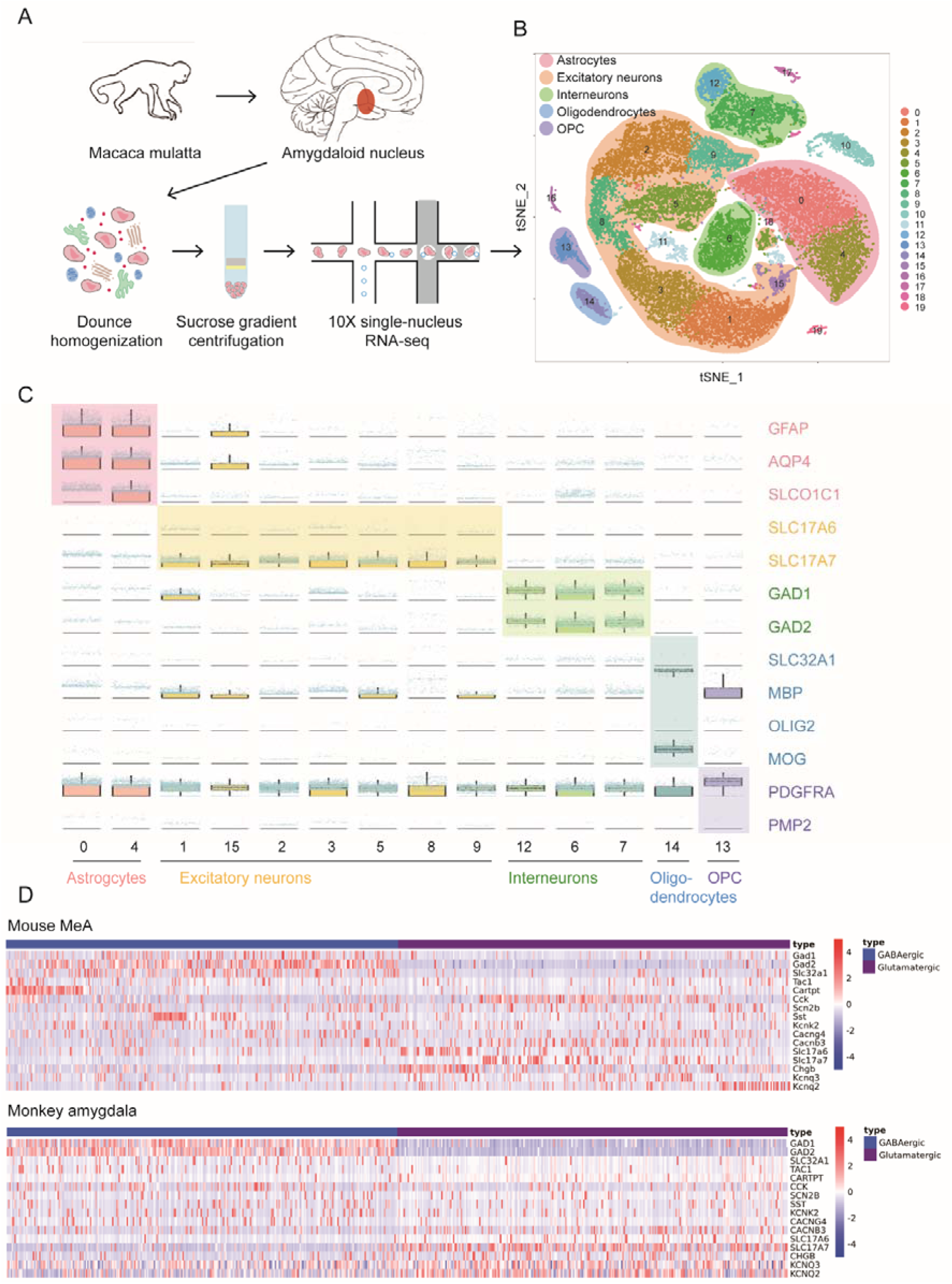
Classification and relative expression of known markers of monkey amygdalae nuclei using single-nucleus RNA-seq. (A) Schematic showing the process of single-nucleus RNA sequencing. The tissues samples of amygdala were surgically removed. The frozen amygdala tissue was homogenized. Then various concentration of sucrose was used to purify the nucleus. The scRNA-Seq libraries were generated using the 10X Genomics Chromium Controller Instrument. (B) Two-dimensional t-distributed stochastic neighbour embedding (tSNE) visualization from single-nucleus RNA sequencing showing the distribution of all nucleus. 20 major classes of nuclei were distinguished. Each dot represents a single nucleus colored according to cluster assignment. The background is colored by major cell types (Astrocytes, Excitatory neurons, GABAergic neurons, Oligodendrocytes, Oligodendrocyte Progenitors cells (OPC)). (C) Boxplots showing the distribution of representative type markers among 14 clusters shown at the bottom. (D) Heatmap showing different expression level of selected top marker genes of GABAergic and Glutamatergic neurons in mouse MeA and monkey amygdala.

We totally obtained 26,400 cells from amygdala of two monkeys for further analysis, in which average 2500 genes and 5500 transcripts were identified in each cell. To classify major cell types in primate amygdala, we performed t-distributed stochastic neighbour embedding (tSNE) analysis using Seurat. We totally identified 19 clusters, among which we further classified them into five major clusters, excitatory neurons, GABAergic neurons, astrocytes, oligodendrocytes and oligodendrocyte progenitor cells (OPCs) (Fig. 1B).

To further analyse sub-clusters within each cell type, we applied the random forest method to segregate 5 major cellular types, according to unique cell-type specific markers (Fig. 1C). Interestingly, in the glutamatergic neurons of primate amygdala, we found that SLC17A7 (VGLUT1) is the major marker gene, instead of SCL17A6 (VGLUT2) which is normally seen in mouse media amygdala (MeA). Whereas for GABAergic neurons, both GAD1 and GAD2 genes are well expressed (Fig. 1C).

In order to understand the evolutionary differences between rodent and primate, we compare expression profiles of major marker genes in the glutamatergic and GABAergic neurons from available datasets^[3, 9]^. The current available single-cell RNA-seq database mainly focus on MeA region of amygdala, where GABAergic neurons are major cell types. Whereas in primate, BLA became the major region in amygdala, thus there are more glutamatergic neurons found in the monkey amygdala snRNA-seq dataset (Fig. 1D)^[2]^.

### Classification of glutamatergic neurons in the primate amygdala

To analyze the sub-cell types within the glutamatergic neurons in the primate amygdala dataset, we further classified 12816 glutamatergic neurons into 19 sub-clusters using tSNE analysis (Fig. 2A). We found that SLC17A7 (VGLUT1) was universally expressed in nearly 19 sub-clusters (Fig. 2B,C). We further performed GO (gene ontology) analysis to examine the differentially expressed gene in glutamatergic neurons (Fig. 2D, Table S2).

**Figure 2.**
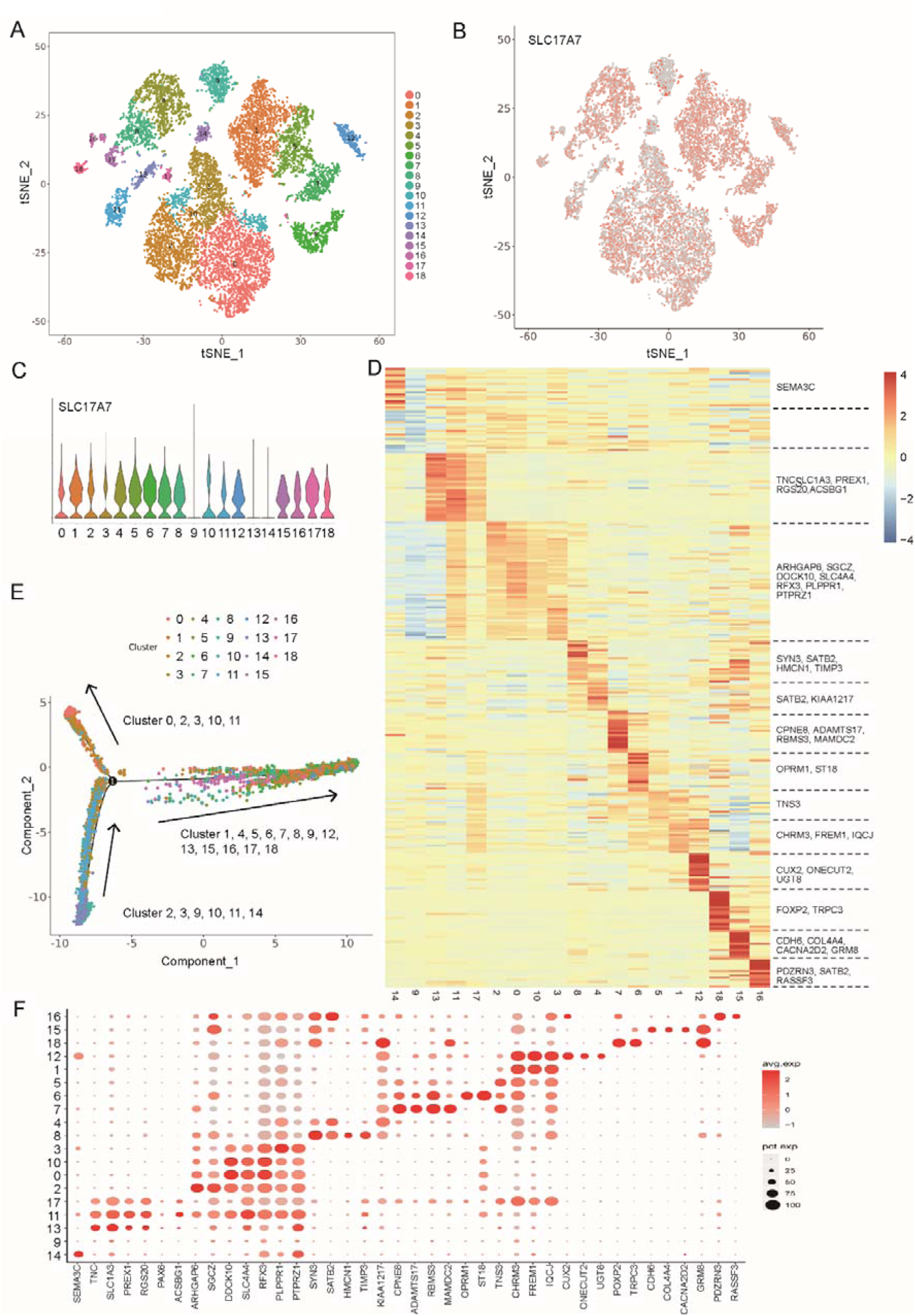
Molecular taxonomy of excitatory neurons in primate amygdala. (A) Visualization of 19 subclusters of excitatory neurons using two-dimensional tSNE. Individual dots correspond to single nucleus colored according to subcluster assignment. (B) Gene expression of known marker SLC17A7 is shown in the tSNE visualization (grey: no expression, red: relative expression). (C) Violin plots showing the distribution of SLC17A7 in different subclusters of excitatory neurons. (D) Heatmap showing the distribution and expression level of significantly differentially expressed genes for each subcluster of excitatory neurons. Major changed genes in each subcluster were listed on the right. (E) Pseudo time showing the possible trajectory of excitatory neurons depended on biomarkers. (F) Expression of representative type-enriched markers among 19 subclusters.

Monocle analysis maps whole-transcriptome profiles of single cells along an artificial temporal curve in orderly fashion to identify the starting and end points of cell trajectories in high-dimensional space^[15, 16]^. We next reconstructed developmental relationships of sub-clusters using Monocle analysis (Fig. 2E). Total 19 sub-clusters exhibited three distinct developmental trajectories, suggesting the different developmental origins and destinations of glutamatergic neurons in the primate amygdala (Fig. 2E).

To further identified the molecular taxonomy of glutamatergic neurons in the primate amygdala, we showed the relative expression level (encode by color depth) and percent of cell expressed (encode by size of circle) of major expressing genes, other than marker genes, in the 19 sub-clusters (Fig. 2F).

### Classification of GABAergic neurons in the primate amygdala

GABAergic neurons play different roles in different regions of the amygdala. For example, GABAergic neurons of the CeA promotes cataplexy in mice and GABAergic neurons of the BLA contribute to post-traumatic stress syndrome (PTSD), autism, attention-deficit hyperactivity disorder (ADHD)^[17, 18]^. To analyse the subtypes within the GABAergic neurons in the primate amygdala dataset, we further classified 4215 GABAergic neurons into 13 sub-clusters using tSNE analysis (Fig. 3A).

**Figure 3.**
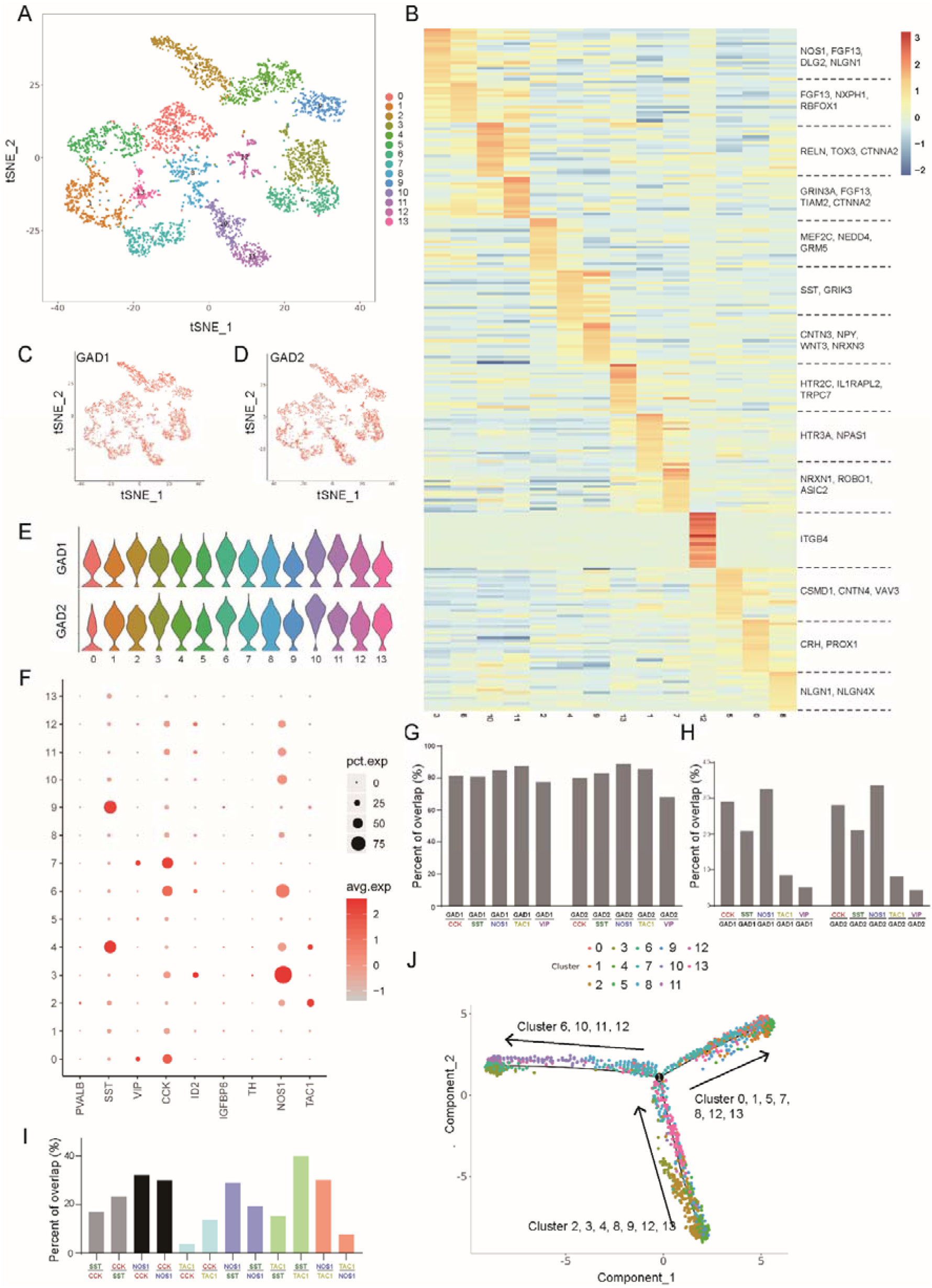
Molecular taxonomy of GABAergic neurons in primate amygdala. (A) Visualization of 14 sub-clusters of GABAergic neurons using two-dimensional tSNE. Individual dots correspond to single nucleus colored according to subcluster assignment. (B) Heatmap showing the distribution and expression level of significantly differentially expressed genes for each sub-cluster of GABAergic neurons. Major changed genes in each sub-cluster were listed on the right. (C-D) Gene expression of known marker GAD1 (C) and GAD2 (D) is shown in the tSNE visualization (grey: no expression, red: relative expression). (E) Violin plots showing the distribution of GAD1 and GAD2 in different subclusters of GABAergic neurons. (F) Expression of representative type-enriched markers among 14 subclusters. (G-I) Quantification of percentage overlap between nuclei labelled by different marker genes. (G) Overlap between primary marker genes for GABAergic neurons GAD1 (or GAD2) and neuropeptide markers (CCK, SST, NOS1, TAC1, VIP). The percentage of overlap was counted as followed, GAD1/CCK representing the percentage of CCK and GAD1 double labelled nuclei in CCK positive nuclei. (H) Reciprocals of (G). (I) Overlap between different neuropeptide markers (CCK, SST, NOS1, TAC1). (J) Pseudo time showing the possible trajectory of GABAergic neurons depended on biomarkers.

To further identified the molecular taxonomy of GABAergic neurons in the primate amygdala, we showed the distribution of expression level of representative subtype GABAergic neurons markers across 13 neuronal sub-clusters, from which there are numerous critical GABAergic cellular markers including NOS1 (nitric oxide synthase 1), RELN, SST (somatostatin), NPY, CCK (cholecystokinin), as well as synaptic molecules such as NLGN1, NRXN3, NRXN1 (Fig. 3B, Table S3). Surprisingly, we found that there were very few parvalbumin-positive neurons in the GABAergic neuron dataset of primate, suggesting that there is a dramatic evolutionary difference from rodent to primate, regarding to the function of parvalbumin-positive neurons in amygdala (Fig. S1).

Both GAD1 and GAD2, two primary marker genes for GABAergic neurons, were highly expressed in all 13 neuronal sub-clusters (Fig. 3C, D, E). The relative expression level (encoded by color depth) and percent of cell expressed (encoded by size of circle) of the subtype marker genes were illustrated in Fig. 3F, from which one could find that SST, CCK, NOS 1, TAC1 (substance P) and VIP (vasoactive intestinal peptide) were five major cellular markers for subtypes of GABAergic neurons in the primate amygdala.

Next, we would like to examine the proportion of each subtype neurons within GABAergic populations. First, we verified the over 80% of SST, CCK, NOS1, TAC1 and VIP-expressing neurons are GAD1 and GAD2-positive neurons (Fig. 3G). From calculation the percentage of each marker in GAD1 or GAD2, it appears that CCK and NOS1-expressing neurons account for 30% of total GABAergic neurons, SST-expressing neurons account for 20%, and TAC1-expressing neurons account for around 10%, whereas VIP-expressing neurons account only for less 5% of GABAergic neurons (Fig. 3H). Across these four major subtypes (CCK, NOS1, SST and TAC1), there are various extent of overlapping (10-40%), as shown in Fig. 3I.

After further analysing differentially expressed genes in these sub-clusters, we found that expression of LHX6 and CUX2, two marker genes for interneuron progenitor cell, were enriched in sub-cluster #2,3,4,9 and #2,4,9 separately. We then constructed developmental relationships of sub-clusters using Monocle analysis. Total 13 sub-clusters exhibited three distinct developmental trajectories, suggesting the different developmental origins and destinations of GABAergic neurons in the primate amygdala (Fig. 3J).

### Subtypes of astroglia and development of oligodendrocytes in primate amygdala

It is speculated that astroglia has underwent enormously expansion in primate species, as the white matter dramatically increased in monkeys and humans comparing to rodent species^[19]^. Thus, it is of great interests to examine the subtypes of astroglia in amygdala of rhesus monkeys.

We classified 5889 astrocytes into 8 clusters using tSNE analysis (Fig. 4A). Previous study classified astrocytes in rodent amygdala three distinct subclasses as AS1, AS2, and AS3^[9]^. Interestingly, by analyzing the differentially expressed gene in astroglia of monkey amygdala we identified two more clusters of astrocytes in the amygdala of primates, which we refer as AS4 (Cluster 7) and AS5 (Cluster 8) (Fig. 4B). Specifically, NPSR1 was selectively enriched in AS4, while T8SIA2 and SEMA3C were selectively enriched in AS5 (Fig. 4C).

**Figure 4.**
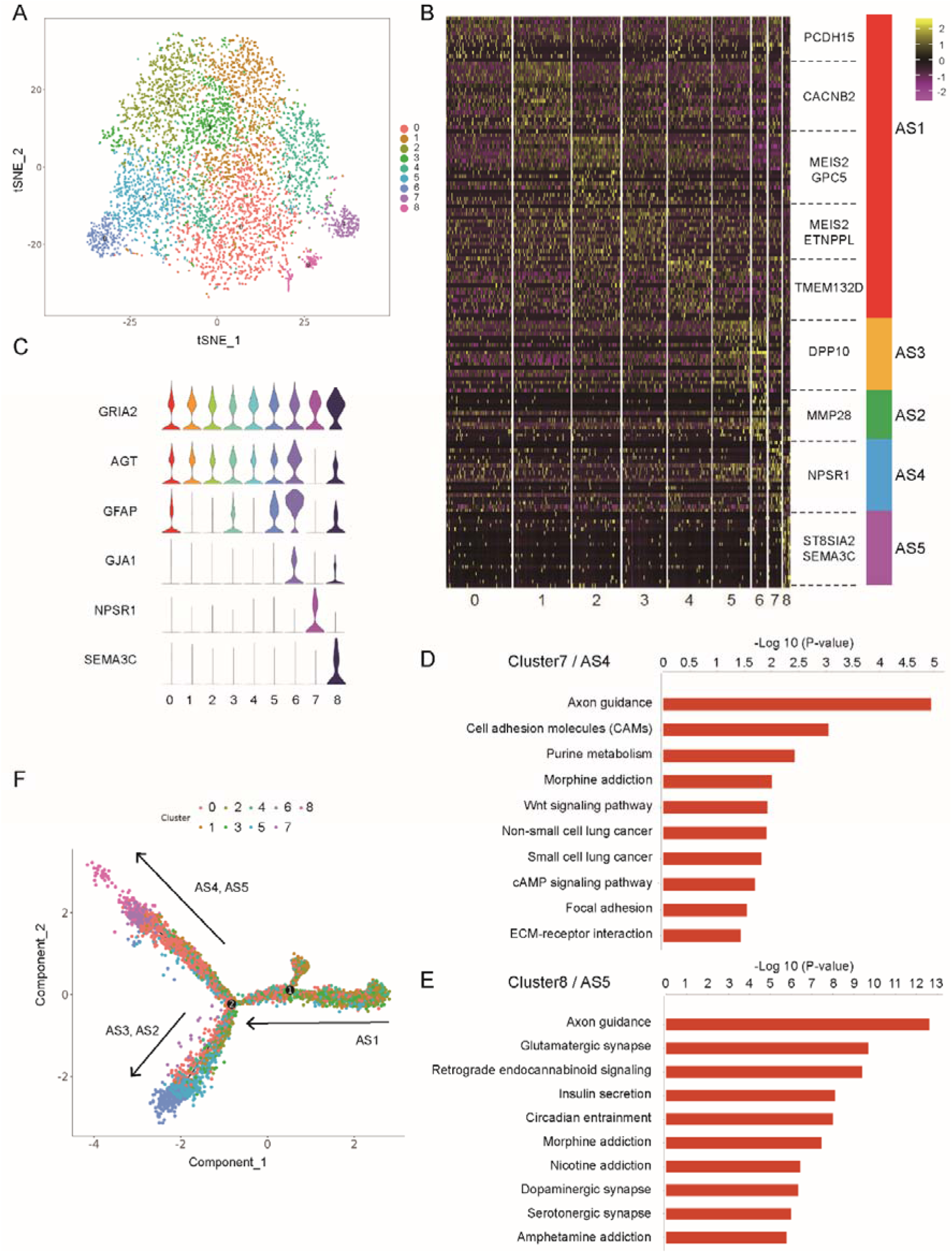
Molecular taxonomy of primate astroglia in amygdala. (A) Visualization of 9 subclusters of astroglia using two-dimensional tSNE. Individual dots correspond to single nucleus colored according to subcluster assignment. (B) Heatmap showing the distribution and expression level of significantly differentially expressed genes for each subcluster of astroglia. Major changed genes in each subcluster were listed on the right. All astroglia were divided as 5 distinct subclasses (AS1∼AS5). Cluster 0 to cluster 4 were generalized into AS1, others were in a one-to-one correspondence. (C) Violin plots showing the distribution of representative type markers among 9 different subclusters of astroglia. (D-E) Top pathway terms enriched in two specific clusters of primates, cluster7/AS4 (D) and cluster8/AS5 (E). (F) Pseudo time showing the possible trajectory of astroglia depended on biomarkers.

Importantly, with performing GO functional analysis we found that gene enriched in both AS4 and AS5 are strongly associated with the axon guidance process, suggesting that the primate-specific astroglia subtypes may contribute the formation of neural circuits (Fig. 4D, E). Furthermore, with monocle analysis we showed that AS1 is an early time point, while AS2, AS3 and AS4, AS5 are two different end points of cell trajectories (Fig. 4F).

In the central nervous system (CNS), the myelin sheaths are formed by oligodendrocytes (OLs) which deafferentation from OPCs. Defects in oligodendrocytes development have been reporter in various neurodegenerative diseases and developmental disorders^[20-22]^. We acquired single cell transcriptomes for 478 OPC, 392 OL and 740 OPC-OL cells, the middle stage between OPC and OL. We classified these cells into 7 sub-clusters using tSNE analysis (Fig. 5A, B). Because PDGFRα, the OPC marker, was enriched in cluster 1 and 6, monocle analysis shown that the group of cells including cluster 1 and 6 was an early time point in a pseudo time alignment (Fig. 5C).

**Figure 5.**
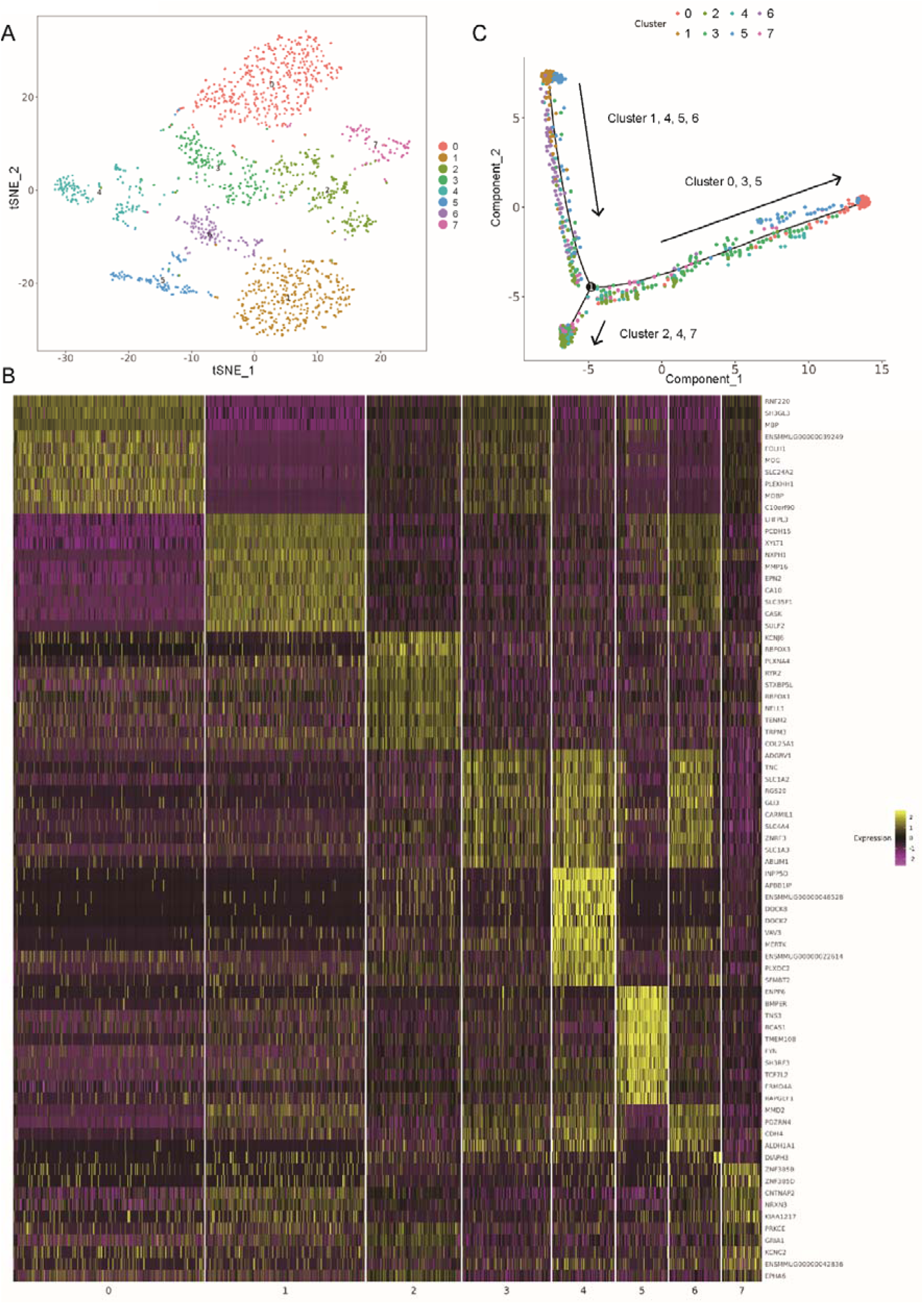
Molecular taxonomy of oligodendrocytes, OPC and OPC-OL cells in primate amygdala. (A) Visualization of 8 subclusters of OPC-OL using two-dimensional tSNE. Individual dots correspond to single nucleus colored according to subcluster assignment. (B) Pseudo time showing the possible trajectory of OPC-OL depended on biomarkers. (C) Heatmap showing the distribution and expression level of significantly differentially expressed genes for each subcluster of OPC-OL.

To further investigate the evolutionary differences between rodent and primate, we finally compare expression profiles of major marker genes in the astrocytes, endothelial, microglia, mural, oligodendrocytes, oligodendrocyte precursor cells from available datasets (Fig. S2A, B)^[3, 9]^.

### Cross-species transcriptome comparison between macaque and mouse amygdala

To investigate the evolutional variation of gene expression profiles between the primate and rodent amygdala, we compared the single cell transcriptomic profiles of macaque dataset and previous mouse dataset^[9]^ using the LIGER algorithm^[23]^ which can recognize the significant shared and dataset-specific gene markers. tSNE visualization of macaque and mouse single cells showed that there are some similarities, but mostly variations between primates and rodents (Fig. 6A, B). The LIGER joint cluster assignments indicated that oligodendrocytes were similar while neurons and astrocytes were rather species-specific, although they were mixed to some extent in the tSNE plot (Fig. 6C). Clusters 2, 3 and 4 were macaque-dominant, yet clusters 0 and 1 were mouse-dominant (Fig. 6C). A gene set in factor 14 showed high expression in cells from cluster 4 and cluster 11, majority of which were NOS1-positive GABAergic neurons (Fig. 6D). The analysis of species-specific genes of factor 14 illustrated that *SPOCK3, FRMD4A* were highly expressed by macaque NOS1-positive GABAergic neurons, whereas *Rlbp1, Olig1* were highly expressed by this type of neurons in mouse (Fig.6D). Although oligodendrocytes from macaque and mouse were similar by LIGER analysis (Fig. 6C), we still found several species-specific genes in the oligodendrocytes related to factor 4, such as genes *ARHGAP22, LRRTM3*, which were specifically expressed in macaque (Fig. 6E).

**Figure 6.**
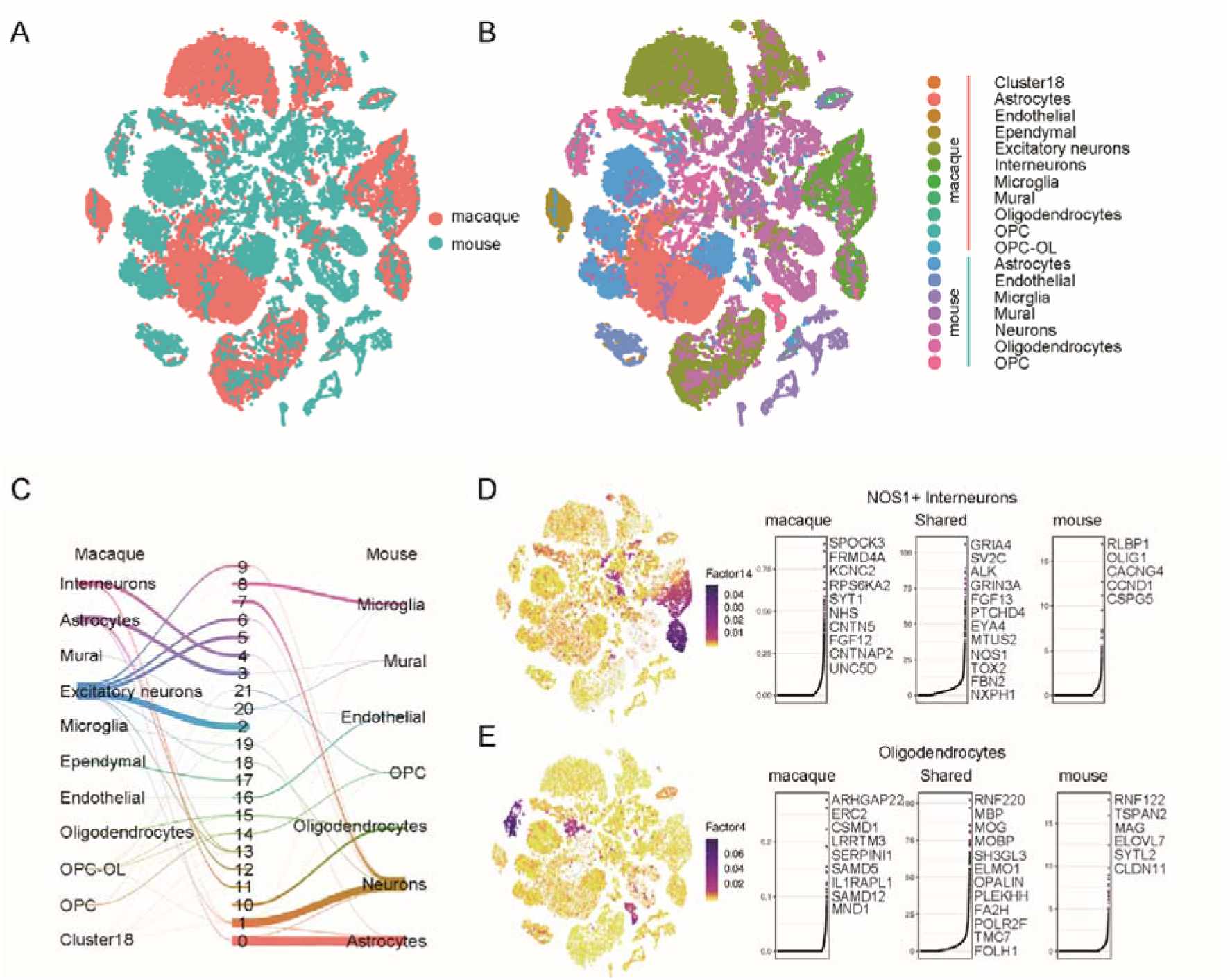
Cross-species transcriptome comparison between macaque and mouse amygdala. (A) tSNE visualization of macaque and mouse single cells analysed by LIGER, color-coded by species. (B) tSNE visualization of macaque and mouse single cells analysed by LIGER, color-coded by cell types. (C) River plots comparing the previously cell types assignments of macaque and mouse with LIGER joint clusters. (D-E) Cell factor loading values (left) and gene loading plots (right) of left loading dataset-specific and shared genes for factor 14(D) and factor 4(E).

## DISCUSSION AND CONCLUSION

Amygdala in mammals is an anatomically complex brain region containing several sub-regions, including medial, central and basal etc. The previous studies for amygdala are mainly focusing on rodents. In rodent medial amygdala play a central role in regulating emotional and other adaptive behaviors, although basal lateral amygdala also showed critical functions. However, in primate the basal lateral amygdala became the primary subregion of amygdala. Thus, it is of essential importance to illustrate the molecular taxonomy and cellular composition of amygdala in primates.

Thanks to the technical advance of single-cell RNA sequencing technology, it is now possible to study gene expression profiles in a single-cell resolution. In our study, we revealed numerous novel cell types in known glutamatergic, GABAergic neuron and astroglia. For example, we found that there are numerous unknown subtypes of glutamatergic neurons in primate amygdala, comparing to rodent amygdala. Furthermore, the composition of GABAergic neurons in primate amygdala are quite different from its of rodent amygdala. In primate amygdala, GAD1 and GAD2 are primary cellular markers for GABAergic neurons and they are co-expressed in majority of GABAergic neurons. Over 80% of the cells expressing five of major neuronal peptide marker genes (CCK, SST, NOS1, TAC1, VIP) are positive for GAD1 and GAD2, which are quite different from rodent, for example only 20% of CCK positive neurons of rodent amygdala are GAD2 positive. Moreover, in primate amygdala, composition of GABAergic neurons (CCK:30%, SST:20%, NOS1:35%, TAC1:10%, VIP:5%) is quite different from rodent amygdala. Moreover, we also found *SPOCK3, FRMD4A* were highly expressed by macaque NOS1-positive GABAergic neurons, whereas *Rlbp1, Olig1* were highly expressed by this type of neurons in mouse. Overall, these differences suggested that the neuronal function of primate amygdala may be distinct from its in rodent.

We discover the largest difference between rodent and primate in cellular composition of astroglia. In rodent, the primary marker genes for astroglia are Gja1, Lfng and Agt, however, we found that the primary marker genes are GRIA2 and AGT in primate amygdala. Furthermore, we found two more novel subcluster of astroglia, AS4 and AS5, which appeared to have unique functions for axon guidance and synapse formation (Fig. 4D). Together, we believe that this dataset is very valuable for our deeper understanding of the neuronal anatomy of primate brain.

## ACKNOWLEDGEMENTS

This work was supported by the National Nature Science Foundation of China (NSFC) Grants (#81970990 to L.Z, #81771132 to L.Z, #81571028 to H.J, #31625013 to Z.L.Q, #91732302 to Z.L.Q), Shanghai Brain-Intelligence Project from STCSM (#16JC1420501 to Z.L.Q), the Strategic Priority Research Program of the Chinese Academy of Sciences (#XDBS01060200 to Z.L.Q) and the Shanghai Municipal Science and Technology Major Project (#2018SHZDZX05 to Z.L.Q). The research is supported by the Open Large Infrastructure Research of Chinese Academy of Sciences. National Key R&D Program of China (#2017YFA0105201 to Z.L.Q) and the CAMS Innovation Fund for Medical Sciences (#CIFMS 2016-I2M-2-001 to Z.L.Q). Shanghai Jiao Tong University School of Medicine Two-hundred Talent-20191818 to L.Z. Foundation of Shanghai Municipal Health Commission (#21840052 to L.Z). Funding for open access charge: National Nature Science Foundation of China.

## Supplementary Materials

**Fig. S1.**
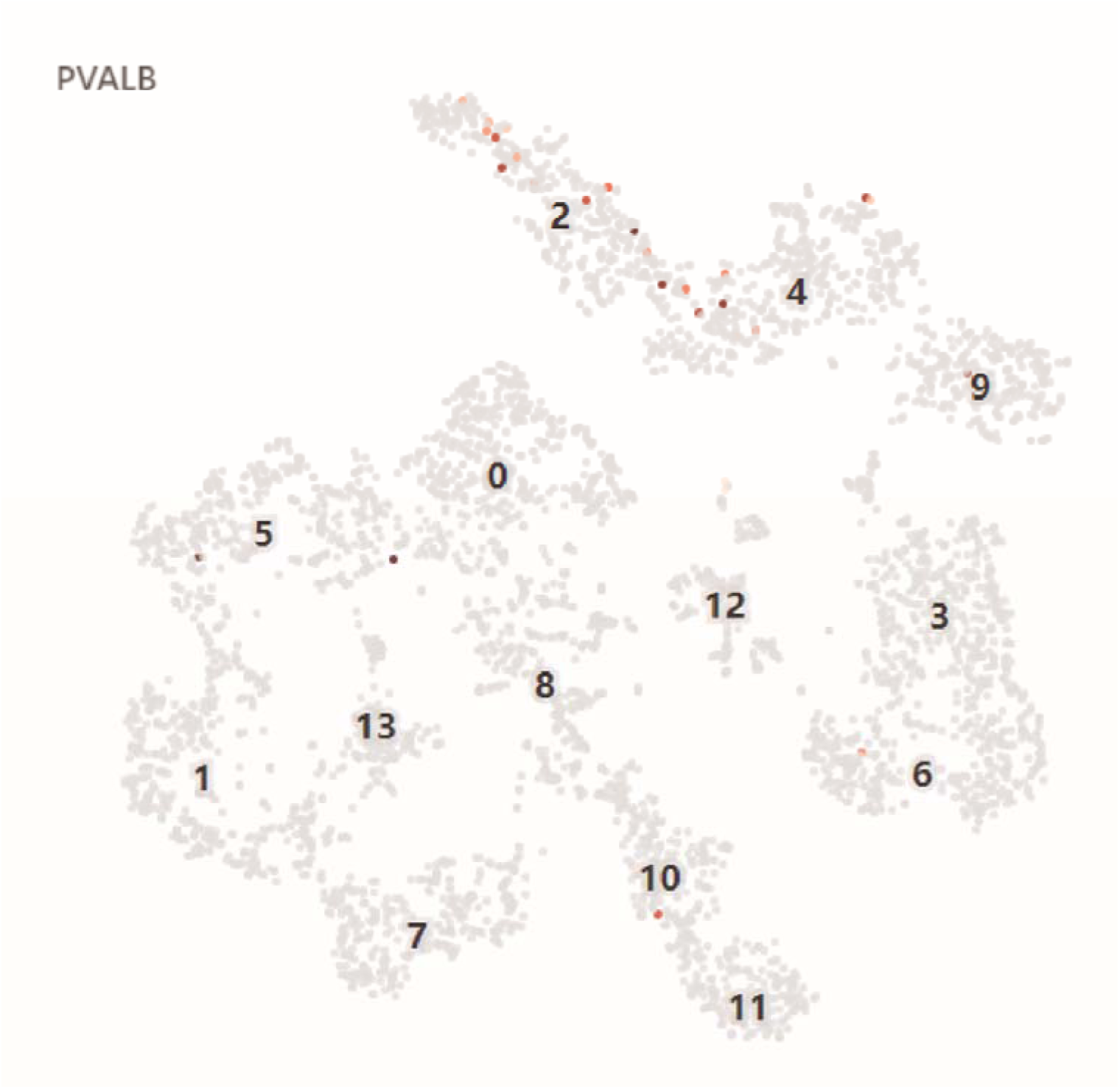
Expression of PVALB in GABAergic neurons of primate amygdala. Gene expression of known marker PVALB is shown in the tSNE visualization of GABAergic neurons (grey: no expression, red: relative expression).

**Fig. S2.**
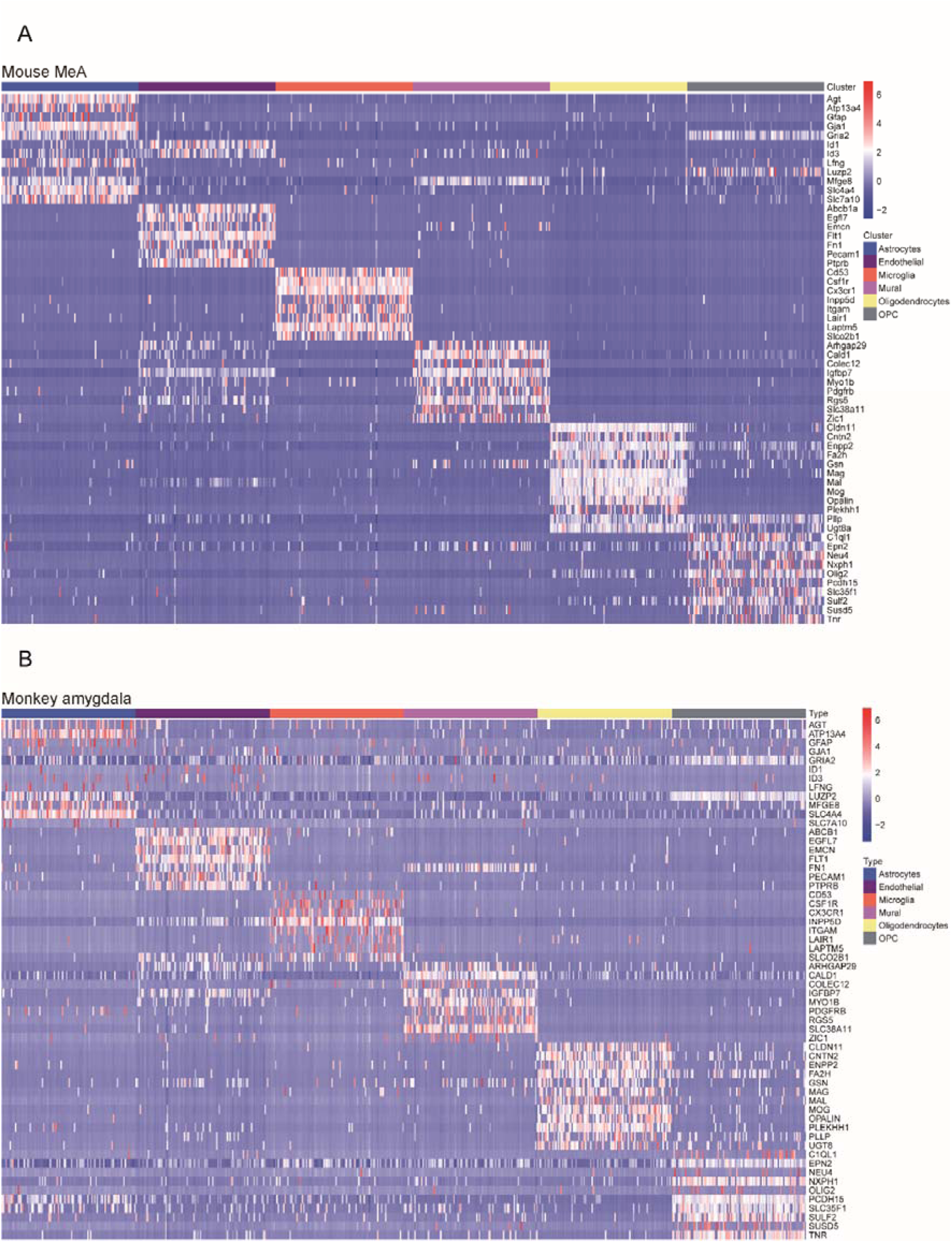
Top marker genes of non-neuronal cells between mouse MeA and monkey amygdala. (A-B) Heatmap showing different expression level of selected top marker genes of Astrocytes, Endothelial, Microglia, Mural, Oligodendrocytes and OPC in mouse MeA (A) and monkey amygdala (B).

**Table S1.**
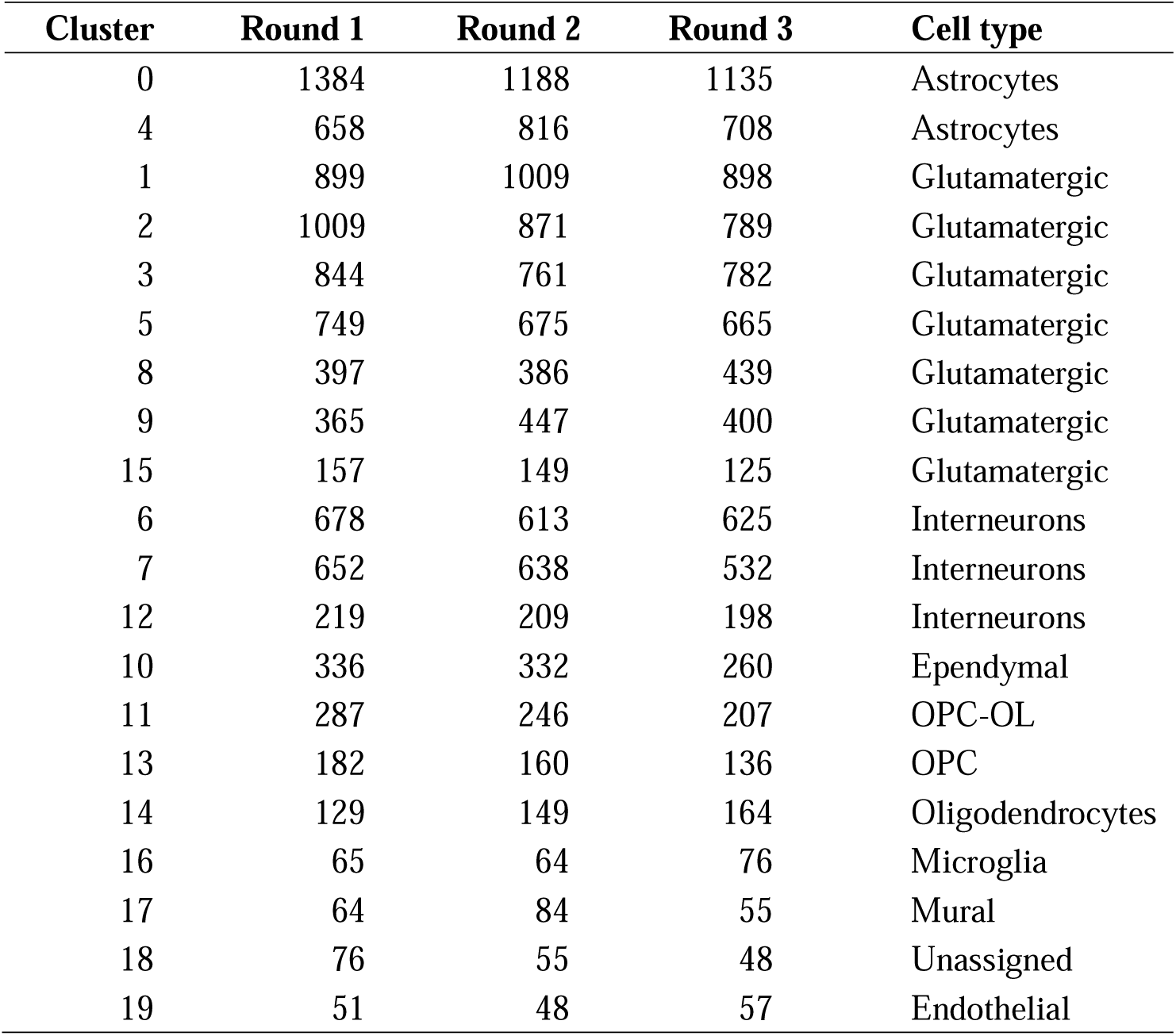
Classification of amygdala cells in rhesus monkey. Clusters were numbered according to cell numbers counted (200 genes, 500 transcripts per cell).

**Table S2. The differentially expressed gene in glutamatergic neurons.** An additional table file shows this in more detail [see Additional file 1-Table S2].

**Table S3. Expression level of representative subtype GABAergic neurons marker genes.** An additional table file shows this in more detail [see Additional file 2-Table S3].

